# CYP4F2 is a human-specific determinant of circulating N-acyl amino acid levels

**DOI:** 10.1101/2023.03.09.531581

**Authors:** Julia T. Tanzo, Veronica L. Li, Amanda L. Wiggenhorn, Maria Dolores Moya-Garzon, Wei Wei, Xuchao Lyu, Wentao Dong, Usman A. Tahir, Zsu-Zsu Chen, Daniel E. Cruz, Shuliang Deng, Xu Shi, Shuning Zheng, Yan Guo, Mario Sims, Monther Abu-Remaileh, James G. Wilson, Robert E. Gerszten, Jonathan Z. Long, Mark D. Benson

## Abstract

N-acyl amino acids are a large family of circulating lipid metabolites that modulate energy expenditure and fat mass in rodents. However, little is known about the regulation and potential cardiometabolic functions of N-acyl amino acids in humans. Here, we analyze the cardiometabolic phenotype associations and genetic regulation of four plasma N-fatty acyl amino acids (N-oleoyl-leucine, N-oleoyl-phenylalanine, N-oleoyl-serine, and N-oleoyl-glycine) in 2,351 individuals from the Jackson Heart Study. N-oleoyl-leucine and N-oleoyl-phenylalanine were positively associated with traits related to energy balance, including body mass index, waist circumference, and subcutaneous adipose tissue. In addition, we identify the *CYP4F2* locus as a human-specific genetic determinant of plasma N-oleoyl-leucine and N-oleoyl-phenylalanine levels. In vitro, CYP4F2-mediated hydroxylation of N-oleoyl-leucine and N-oleoyl-phenylalanine results in metabolic diversification and production of many previously unknown lipid metabolites with varying characteristics of the fatty acid tail group, including several that structurally resemble fatty acid hydroxy fatty acids (FAHFAs). By contrast, FAAH-regulated N-oleoyl-glycine and N-oleoyl-serine were inversely associated with traits related to glucose and lipid homeostasis. These data uncover a human-specific enzymatic node for the metabolism of a subset of N-fatty acyl amino acids and establish a framework for understanding the cardiometabolic roles of individual N-fatty acyl amino acids in humans.

## Main

The N-acyl amino acids are a large and structurally diverse family of circulating lipid signaling molecules. These lipid metabolites are unusual peptide conjugates of fatty acids and amino acids. In mice, levels of specific circulating N-acyl amino acids are tightly controlled by the action of two enzymes, PM20D1 and FAAH, which catalyze bidirectional N-acyl amino acid synthesis from and hydrolysis to free fatty acids and free amino acids^1–4^. Pharmacological, genetic, and mechanistic studies in rodents suggest that certain members of the N-acyl amino acids stimulate mitochondrial respiration and whole-body energy expenditure^1^. Other complementary studies have also established roles for N-acyl amino acids in glucose homeostasis^2^, adipogenesis^5^, vascular tone^6, 7^, and bone homeostasis^8, 9^. Importantly, the functional consequence and enzymatic regulation of each N-acyl amino acid is highly dependent on the structural properties of the fatty acid tail group and amino acid head group. For example, even subtle modification of the lipid chain or amino acid moieties can effect N-acyl-amino acid substrate specificity for PM20D1 and ligand potency in mitochondrial respiration^1^.

Despite the considerable body of rodent literature on N-acyl amino acids, our knowledge of the regulation and clinical associations of these lipid metabolites in humans is still lacking. Two small studies have previously investigated the role of PM20D1 in the regulation of N-acyl amino acids in human plasma. In one report, plasma N-oleoyl-leucine and N-oleoyl-phenylalanine levels were found to be associated with circulating PM20D1 protein levels and correlated positively with adiposity and several parameters of glucose homeostasis^10^. However, a second study failed to identify any association of serum levels of two N-acyl amino acids, N-oleoyl-leucine or N-oleoyl-glutamine, with genetic variants in the *PM20D1* locus though in a very small cohort^11^. The conflicting human data, as well as the scope of phenotypes and limited number of individuals examined in the previous two studies therefore motivate additional investigations as to the clinical relevance and genetic underpinnings of this class of molecules.

Our group recently measured circulating levels of four N-acyl-amino acids (N-oleoyl-glycine, N-oleoyl-serine, N-oleoyl-leucine, and N-oleoyl-phenylalanine) in plasma samples from 2,351 participants of the Jackson Heart Study (JHS) using liquid chromatography-mass spectrometry (LC-MS)^12^. In two broad metabolomic-wide association studies in this population, we identified that levels of N-oleoyl serine and N-oleoyl leucine were among the top metabolites in human plasma associated with future risk of coronary heart disease^13^ and heart failure^12^, respectively. These observations extend prior studies in murine models and suggest that additional investigations of N-acyl amino acid biology may provide important new insights into human metabolism and cardiometabolic disease.

Here, we provide a detailed analysis of the role of the oleic fatty acid tail group and amino acid head group in driving associations with cardiometabolic disease. We use genome wide association studies (GWAS) to distinguish between different molecular species of N-acyl amino acids according to amino acid head group. Further, this analysis identifies a previously unknown enzymatic regulator of N-oleoyl-leucine and N-oleoyl-phenylalanine, CYP4F2, in humans. Finally, we use untargeted metabolomics in live cells to map the fate of N-oleoyl-leucine and N-oleoyl-phenylalanine downstream of CYP4F2 hydroxylation and identify numerous previously unknown lipid metabolites with varying characteristics of the fatty acid tail group, including several that structurally resemble fatty acid hydroxy fatty acids (FAHFAs), a recently-discovered family of bioactive lipids that signal through specific G protein-coupled receptors to improve glucose-insulin homeostasis and block inflammatory cytokine production^14^. Our data integrating mass spectrometry, human phenotyping, genetics, and model systems provide new insights to this emerging class of molecules.

## Results

### N-acyl amino acids are associated with cardiometabolic disease independent of free amino acid plasma levels and according to the amino acid head group

We recently measured circulating levels of N-oleoyl-glycine, N-oleoyl-serine, N-oleoyl-leucine, and N-oleoyl-phenylalanine in plasma samples from 2,351 participants of the JHS using liquid chromatography-mass spectrometry (LC-MS), as described^12^. Baseline characteristics of the JHS study participants are detailed in **Supplemental Table 1**. Age- and sex-adjusted analyses identified N-oleoyl-serine and N-oleoyl-leucine as top metabolites associated with the future risk of coronary heart disease^13^ and heart failure^12^, respectively. However, prior work did not examine whether the associations of the intact N-acyl oleic acid conjugate of each amino acid were independent of the corresponding free amino acid (**Fig. 1a**). Using age-, sex-, and batch-adjusted Cox regression models, we determined that the association between N-oleoyl-serine and the future risk of coronary heart disease remained unaffected when further adjusted for free serine (hazard ratio [HR] 0.81 per 1 SD increment in transformed and normalized metabolite level; 95% CI, 0.69 to 0.94; p-value = 7.3×10^-3^; median follow up 11.6 years). Similarly, the association between N-oleoyl-leucine and future heart failure remained highly significant when further adjusted for free leucine (HR 0.78; 95% CI, 0.66 to 0.91; p-value = 2.1×10^-3^; median follow up 12.6 years). These observations may point towards additional biological mechanisms conferred by the oleic acid tail group of N-acyl serine and N-acyl glycine.

**Fig. 1.**
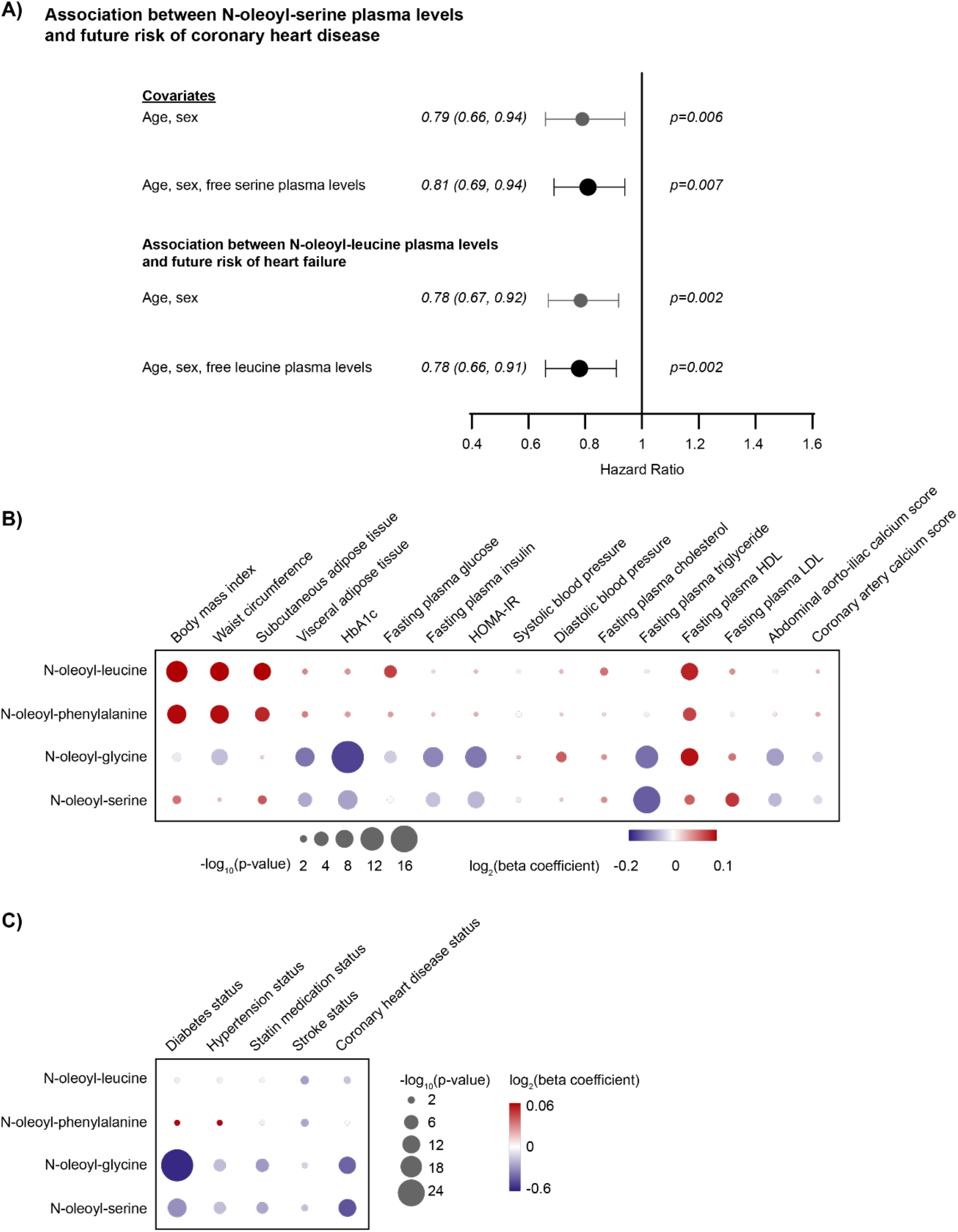
N-acyl amino acids are associated with cardiometabolic disease independent of free amino acid plasma levels and according to the amino acid head group. (A) Age- and sex-adjusted associations between plasma levels of N-oleoyl-serine and future risk of coronary heart disease^13^ (top) and N-oleoyl-leucine and future risk of heart failure^12^ (bottom) in JHS were unaffected by further adjustment for circulating levels of the corresponding free amino acid using cox regression models. (B) The relationship between plasma levels of each N-acyl amino acid (log transformed and standardized) and cardiometabolic traits (standardized) was analyzed using age- and sex-adjusted mixed linear regression models for continuous variables measured in the JHS (top) and age- and sex-adjusted logistic regression models for categorical variables in the JHS (bottom). Estimated beta coefficients are represented by a color scale from red to blue (red represents positive and blue represents negative association). The area of each node corresponds to -log10(p-values). Bonferroni-adjusted P-value = 6.0×10^-4^ (0.05/21 tested traits/4 analytes).

We next investigated how the amino acid head group of each N-acyl amino acid contributed to associations with cardiometabolic disease. As shown in **Fig. 1b**, we found that the associations of measured N-acyl amino acids with cardiometabolic traits in JHS bifurcated into two classes depending on the characteristics of the head group. Whereas N-oleoyl-glycine and serine were associated with “protective” traits, showing strong inverse associations with prevalent diabetes (N-oleoyl-glycine beta = -0.58, p-value = 3.5×10^-25^; N-oleoyl-serine beta = – 0.30, p-value = 3.1×10^-9^), prevalent hypertension (N-oleoyl-glycine beta = -0.18, p-value = 2.3×10^-4^; N-oleoyl-serine beta = -0.17, p-value = 2.7×10^-4^), and prevalent coronary heart disease (N-oleoyl-glycine beta = -0.42, p-value = 5.2×10^-7^; N-oleoyl-serine beta = -0.46, p-value = 8.1×10^-8^), N-oleoyl-leucine and phenylalanine demonstrated positive associations with cardiometabolically “disadvantageous” traits, such as body mass index (N-oleoyl-leucine beta = 0.12, p-value = 1.5×10^-8^; N-oleoyl-phenylalanine beta = 0.10, p-value = 1.9×10^-6^), waist circumference (N-oleoyl-leucine beta = 0.10, p-value = 2.5×10^-6^; N-oleoyl-phenylalanine beta = 0.09, p-value = 5.6×10^-6^), and subcutaneous adipose tissue (N-oleoyl-leucine beta = 0.10, p-value = 5.6×10^-5^; N-oleoyl-phenylalanine beta = 0.08, p-value = 6.1×10^-4^) (**Fig. 1b**). The inverse associations between N-oleoyl-glycine and serine with cardiometabolic disease remained highly significant when further adjusted for BMI (**Supplemental Table 2**).

### Genetic loci associated with levels of N-acyl amino acids in human plasma

In order to determine if the genetic determinants of N-acyl amino acid plasma levels also differ according to amino acid head group, we leveraged available whole genome sequencing data (NHGRI-EBI GWAS Catalog study GCP000239^15^) in the JHS to compare genome wide associations between oleic acid N-acyl conjugates of serine, glycine, leucine, and phenylalanine, as described^16^. As shown in **Fig. 2A**, we identified a strong association between the fatty acid amide hydrolase (*FAAH*) genetic locus and plasma levels of N-oleoyl-glycine (sentinel variant rs324420, p-value = 5.7×10^-64^, beta 0.54, n=2463) and N-oleoyl-serine (sentinel variant rs324420, p-value = 3.6×10^-32^, beta 0.38, n=2101). These findings are consistent with studies in cell- and murine-based systems that have identified FAAH as an intracellular N-acyl amino acid synthase/hydrolase. Interestingly, the sentinel variant for both metabolites was a common (observed MAF = 0.21 in JHS and global MAF 0.21 as reported in the NCBI Allele Frequency Aggregator), missense variant (chr1:46405089C>A; Pro129Thr) that has previously been shown to reduce FAAH stability and enzymatic activity in cell-based model systems and has been associated with overweight and obesity in human studies^17, 18^.

**Fig. 2.**
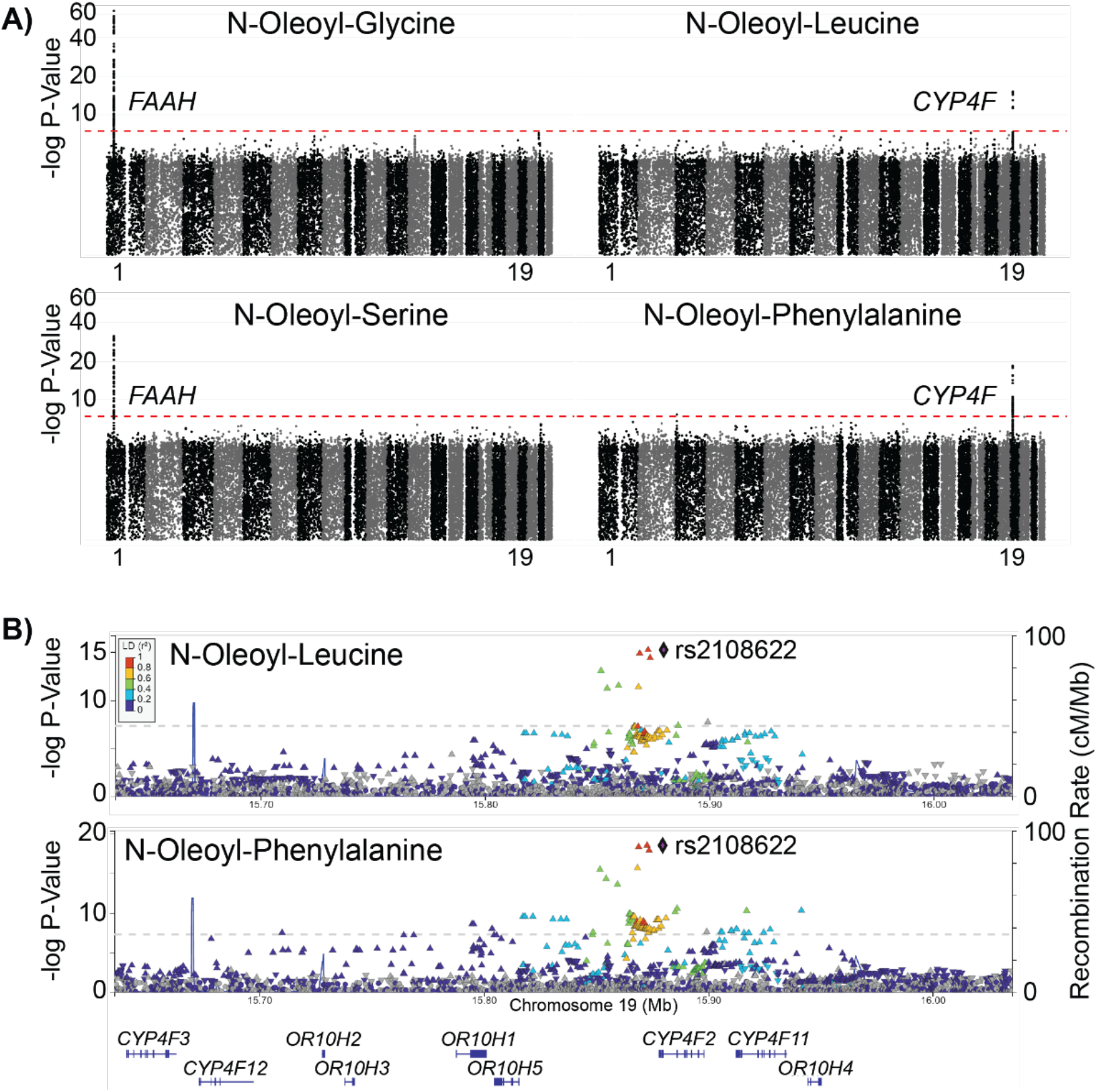
Whole genome association studies of N-acyl amino acid levels in human plasma. (A) Circulating levels of N-oleoyl-glycine and serine in plasma samples from participants from the Jackson Heart Study were associated with the Fatty Acid Amide Hydrolase (*FAAH*) locus on chromosome 1, whereas levels of N-oleoyl-leucine and N-oleoyl-phenylalanine were associated with the Cytochrome P450 Family 4 Subfamily F Member 2 (*CYP4F2*) locus on chromosome 19. Manhattan plots represent the location of each genetic variant across each chromosome (alternating colors; chromosomes 1 and 19 labeled) along the x-axis with the statistical significance of association with the measured metabolite on the y-axis. The genome-wide level of statistical significance (p=5×10^-8^) is marked with a red dashed line. (B) Locuszoom plots of the association signals in *CYP4F2* for N-oleoyl-leucine and N-oleoyl-phenylalanine with the sentinel variant labeled (rs2108622, chr19:15879621C>T; Val433Met).

However, we were interested to find that the *FAAH* locus was not associated with plasma levels of N-oleoyl-leucine or N-oleoyl-phenylalanine in humans at genome-wide (p-value ≤ 5 x 10^-8^), or even nominal levels of statistical significance (sentinel variant rs324420 association with N-oleoyl-leucine p-value = 0.29 and N-oleoyl-phenylalanine p-value = 0.13). Additionally, all four N-acyl amino acids demonstrated no significant association with the *PM20D1* locus (p-values ≥ 0.001 for the associations between each N-acyl amino acid and all variants between the *PM20D1* transcriptional start and end sites, chr1:205,828,025-205,850,132). Instead, we unexpectedly identified a strong association between the *CYP4F2* locus on chromosome 19 and N-oleoyl-leucine (sentinel variant rs2108622, p-value = 5.7×10^-16^, beta 0.44, n = 2101) and N-oleoyl-phenylalanine (sentinel variant rs2108622, p-value = 2.9×10^-19^, beta 0.47, n = 2099) (**Fig. 2B**). CYP4F2 is a cytochrome P450 that catalyzes the omega-hydroxylation of lipophilic substrates.

Taken together, these human genetic data point to specific pathways that regulate circulating N-acyl amino acid levels according to the amino acid head group. In particular, the association of the *CYP4F2* locus with circulating N-oleoyl-leucine and N-oleoyl-phenylalanine was unexpected, and the role of CYP4F2 in N-acyl amino acid metabolism has not been explored.

### CYP4F2-mediated omega-hydroxylation of N-oleoyl-leucine and N-oleoyl-phenylalanine

Canonically, CYP4F2 catalyzes the omega (e.g., terminal) hydroxylation of specific lipophilic substrates, including fatty acids (e.g., arachidonic acid)^19^ and vitamins (e.g., vitamin E and K)^20, 21^. The identification of CYP4F2 as a potential enzymatic regulator of N-oleoyl-leucine/phenylalanine was surprising since this enzyme had not been previously implicated in the metabolism of fatty acid-amino acid conjugates. Nevertheless, we reasoned that CYP4F2 might also catalyze the omega hydroxylation of N-oleoyl-leucine and N-oleoyl-phenylalanine (**Fig. 3A**). To directly test this hypothesis *in vitro*, we first generated lysates overexpressing CYP4F2 by transfection of plasmids expressing human *CYP4F2* into HEK293T cells (**Fig. 3b, left**). Transfected whole cell lysates were then assayed for the ability to catalyze N-oleoyl-leucine and N-oleoyl-phenylalanine hydroxylation in the presence of NADPH. As shown in **Fig. 3B** **(right)**, CYP4F2-transfected cell lysates exhibited a robust hydroxylation activity on both N-acyl amino acids, leading to production of two previously unknown lipid species *in vitro*, hydroxy-oleoyl-leucine and hydroxy-oleoyl-phenylalanine. The absolute rates of CYP4F2-mediated hydroxylation of these N-acyl amino acids were similar to that of a canonical CYP4F2 substrate, arachidonic acid (**Supplementary Fig. 1A**). Additionally, we observed increasing formation rates of hydroxy-oleoyl-leucine and hydroxy-oleoyl-phenylalanine across a range of N-oleoyl-leucine/phenylalanine substrate concentrations (Fig. 3C). Notably, this range of low-mid μM concentrations of N-oleoyl-leucine/phenylalanine correlate to prior studies that detected functional effects of these N-acyl amino acids on energy homeostasis at similar murine model plasma levels and cell-culture treatment doses. The use of CYP4F2-transfected lysates precluded the ability to calculate the catalytic parameters, selectivity, or specificity of this reaction. These data demonstrate that CYP4F2 catalyzes the hydroxylation of N-oleoyl-leucine and N-oleoyl-phenylalanine *in vitro* and provide a biochemical explanation for the observed association between the *CYP4F2* gene locus and circulating N-oleoyl-leucine and N-oleoyl-phenylalanine levels in humans.

**Fig. 3.**
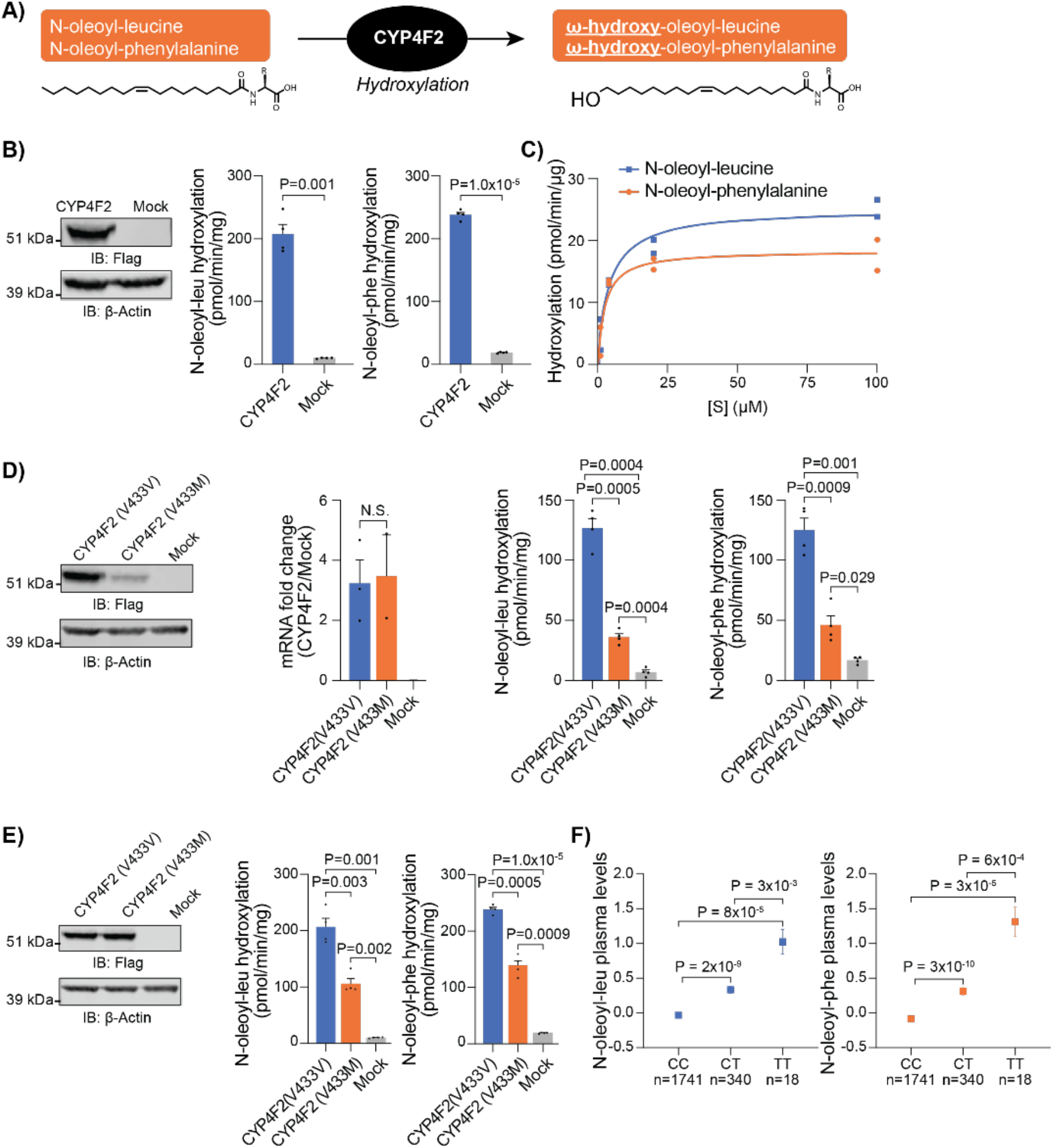
N-oleoyl-leucine and N-oleoyl-phenylalanine are substrates of CYP4F2. (A) Schematic of CYP4F2-mediated hydroxylation of N-oleoyl-leucine and N-oleoyl-phenylalanine. (B) N-oleoyl-leucine and N-oleoyl-phenylalanine hydroxylation in cell lysates of CYP4F2-flag and mock transfected HEK293T cells (N=4) and anti-Flag or anti-β-Actin Western blotting of cell lysates. (C) Dose response analyses of N-oleoyl-leucine and N-oleoyl-phenylalanine hydroxylation by CYP4F2 transfected HEK293T cell lysates (N=2). (D) N-oleoyl-leucine and N-oleoyl-phenylalanine hydroxylation in cell lysates of CYP4F2(V433V)-flag, CYP4F2(V433M)-flag, and mock transfected HEK293T cells (N=4). Cells were transfected with equal amounts of mRNA and Western blotting of cell lysates was performed with anti-Flag or anti-β-Actin. (E) N-oleoyl-leucine and N-oleoyl-phenylalanine hydroxylation in cell lysates of CYP4F2(V433V)-flag, CYP4F2(V433M)-flag, and mock transfected HEK293T cells (N=4) and anti-Flag or anti-β-Actin Western blotting of cell lysates showing normalized levels of CYP4F2(V433V) and CYP4F2(V433M) enzymes. (F) Participants in the JHS were categorized according to carriage of the common CYP4F2 Val433Met (chr19:15879621C>T) missense variant. Holm-Bonferroni adjusted P-values were generated from age- and sex-adjusted regression analyses on log-transformed and standardized normalized metabolite levels. Data are shown as mean ± SEM.

The sentinel variant in the *CYP4F2* locus that was associated with N-oleoyl-leucine and N-oleoyl-phenylalanine plasma levels in our human studies was a chr19:15879621C>T missense variant that results in a Val433Met substitution (**Fig. 2B**). To examine the consequences of the specific CYP4F2 (V433M) variant on N-oleoyl-leucine and N-oleoyl-phenylalanine hydroxylation activity, we performed similar *in vitro* hydroxylation activity assays with CYP4F2 (V433V) or CYP4F2 (V433M) enzymes. Equivalent transfection for both alleles was confirmed by equivalent mRNA levels (**Fig. 3D**). Under these conditions, both the CYP4F2 protein as well as N-oleoyl-leucine, N-oleoyl-phenylalanine, and arachidonic acid hydroxylation activities of CYP4F2(V433M) were lower than that of CYP4F2(V433V) (**Fig. 3D** **and Supplementary Fig. 1B**). To determine the relative activity of CYP4F2 (V433V) or CYP4F2 (V433M) under conditions of equal protein loading, we normalized levels of CYP4F2 (V433M) and CYP4F2 (V433V) protein as assessed by Western blotting (**Fig. 3E**). CYP4F2 (V433M) exhibited reduced hydroxylation of N-oleoyl-leucine, N-oleoyl-phenylalanine, and arachidonic acid compared to CYP4F2 (V433V) (**Fig. 3E** **and Supplementary Fig. 1C**). These data demonstrate that CYP4F2 (V433M) has both lower catalytic efficiency as well as reduced protein levels relative to CYP4F2 (V433V).

To further characterize the functional consequence of the CYP4F2 on circulating levels of N-acyl amino acids in human plasma, we analyzed levels of N-oleoyl-leucine and N-oleoyl-phenylalanine in participants of the JHS categorized according to carriage of the common CYP4F2 Val433Met (chr19:15879621C>T) missense variant. As shown in **Fig. 3F**, we identified that heterozygous carriers of the *CYP4F2*(19:15879621C>T) genotype in the JHS (N=340) had higher circulating levels of plasma N-oleoyl-leucine (log-transformed and standardized level = 0.3, p-value 1.9×10^-9^) and N-oleoyl-phenylalanine (log-transformed and standardized level = 0.3, p-value 3.3×10^-10^) compared to non-carriers (log-transformed and standardized level of N-oleoyl-leucine = 0.0; N-oleoyl-phenylalanine = -0.1; N=1741). Similarly, homozygote carriers of the *CYP4F2* (19:15879621C>T) genotype (N=18) had even higher plasma levels of N-oleoyl-leucine (log-transformed and standardized level = 1.02, p-value 3.3×10^-3^) and N-oleoyl-phenylalanine (log-transformed and standardized level = 1.31, p-value 5.6×10^-4^) compared to heterozygotes. We therefore conclude that a common missense variant in *CYP4F2* leading to elevated plasma N-oleoyl-leucine and N-oleoyl-phenylalanine in humans also reduces CYP4F2 N-oleoyl-leucine and N-oleoyl-phenylalanine hydroxylation activity *in vitro*.

### N-oleoyl-leucine and N-oleoyl-phenylalanine are competitive inhibitors of CYP4F2

The ability of CYP4F2 to hydroxylate multiple endogenous lipid substrates raises the possibility that active site competition might modulate the relative flux of each substrate through CYP4F2 (**Fig. 4A**). To mechanistically examine whether such substrate competition might occur, we used our *in vitro* CYP4F2 enzyme activity assay to directly measure substrate competition between N-oleoyl-leucine and N-oleoyl-phenylalanine on the canonical CYP4F2 substrate arachidonic acid. At a 1:1 molar ratio, N-oleoyl-leucine efficiently inhibited CYP4F2-mediated conversion of arachidonic acid to 20-HETE (86% suppression, **Fig. 4B**). Similar results were observed with N-oleoyl-phenylalanine (76% suppression, **Fig. 4B**). A dose response curve demonstrated that either N-oleoyl-leucine or N-oleoyl-phenylalanine could inhibit CYP4F2-mediated arachidonic acid hydroxylation even at substoichiometric levels (10 mol%, **Fig. 4B**). To examine the potential generality of N-oleoyl-leucine and N-oleoyl-phenylalanine inhibition of CYP4F2, we performed similar in vitro competition experiments with two additional CYP4F2 substrates, docosanoic acid and 8-HETE. As shown in **Fig. 4C-D**, both N-oleoyl-leucine and N-oleoyl-phenylalanine competed CYP4F2-mediated hydroxylation of both 8-HETE and docosanoic acid, but with differences in potency. For instance, even at substoichiometric levels (10 mol %), both N-oleoyl-leucine and N-oleoyl-phenylalanine strongly suppressed CYP4F2-mediated docosanoic acid hydroxylation (87% inhibition by N-oleoyl-leucine and 81% inhibition by N-oleoyl-phenylalanine, **Fig. 4C**). In contrast, superstoichiometric levels (10-fold excess) of either N-acyl amino acid was required to observe similar levels of competition of 8-HETE hydroxylation (94% inhibition by N-oleoyl-leucine and 84% inhibition by N-oleoyl-phenylalanine, **Fig. 4D**).

**Fig. 4.**
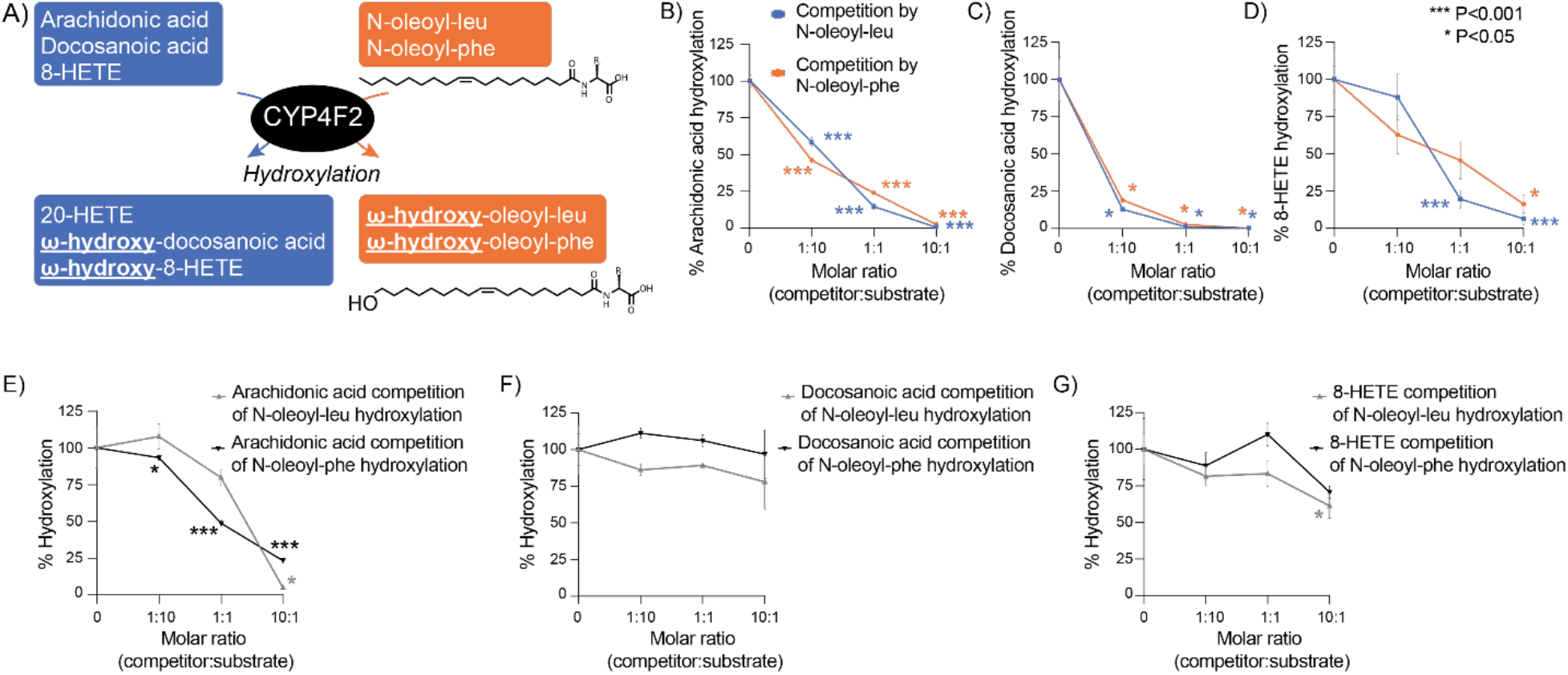
N-oleoyl-leucine and N-oleoyl-phenylalanine are also competitive inhibitors of CYP4F2. (A) Schematic of CYP4F2-mediated hydroxylation of known substrates, N-oleoyl-leucine, and N-oleoyl-phenylalanine. (B) Inhibition of CYP4F2-mediated arachidonic acid hydroxylation by N-oleoyl-leucine and N-oleoyl-phenylalanine in CYP4F2 transfected HEK293T cell lysates (N=4). Cell lysates were treated with indicated molar ratios of N-oleoyl-leucine/phenylalanine and arachidonic acid. Hydroxylation activity of mock transfected HEK293T lysates was subtracted from each point. (C) Inhibition of CYP4F2-mediated docosanoic acid hydroxylation by N-oleoyl-leucine and N-oleoyl-phenylalanine in CYP4F2 transfected HEK293T cell lysates (N=3). Cell lysates were treated with indicated molar ratios of N-oleoyl-leucine/phenylalanine and docosanoic acid. Hydroxylation activity of mock transfected HEK293T lysates was subtracted from each point. (D) Inhibition of CYP4F2-mediated 8-HETE hydroxylation by N-oleoyl-leucine and N-oleoyl-phenylalanine in CYP4F2 transfected HEK293T cell lysates (N=3-4). Cell lysates were treated with indicated molar ratios of N-oleoyl-leucine/phenylalanine and 8-HETE. Hydroxylation activity of mock transfected HEK293T lysates was subtracted from each point. (E) Inhibition of CYP4F2-mediated N-oleoyl-leucine and N-oleoyl-phenylalanine hydroxylation by arachidonic acid in CYP4F2 transfected HEK293T cell lysates (N=4). Cell lysates were treated with indicated molar ratios of arachidonic acid and N-oleoyl-leucine/phenylalanine. Hydroxylation activity of mock transfected HEK293T lysates was subtracted from each point. (F) Inhibition of CYP4F2-mediated N-oleoyl-leucine and N-oleoyl-phenylalanine hydroxylation by docosanoic acid in CYP4F2 transfected HEK293T cell lysates (N=2-3). Cell lysates were treated with indicated molar ratios of docosanoic acid and N-oleoyl-leucine/phenylalanine. Hydroxylation activity of mock transfected HEK293T lysates was subtracted from each point. (G) Inhibition of CYP4F2-mediated N-oleoyl-leucine and N-oleoyl-phenylalanine hydroxylation by 8-HETE in CYP4F2 transfected HEK293T cell lysates (N=3). Cell lysates were treated with indicated molar ratios of 8-HETE and N-oleoyl-leucine/phenylalanine. Hydroxylation activity of mock transfected HEK293T lysates was subtracted from each point.

Conversely, we investigated if the canonical substrates arachidonic acid, docosanoic acid, and 8-HETE might also compete CYP4F2-catalyzed N-acyl amino acid hydroxylation. While arachidonic acid was able to compete N-oleoyl-leucine and N-oleoyl-phenylalanine hydroxylation by CYP4F2 (**Fig. 4E**), neither docosanoic acid nor 8-HETE exhibited strong inhibition of either N-oleoyl-leucine or N-oleoyl-phenylalanine, even at superstoichiometric levels (10:1 competitor:substrate, **Fig. 4F-G**). We therefore conclude that N-oleoyl-leucine and N-oleoyl-phenylalanine and other canonical substrates engage in bidirectional competition to regulate hydroxylation flux at the CYP4F2 enzyme node. The different dose responses observed for competition between each substrate pair potentially reflects differences in effective concentrations and/or substrate affinities at the CYP4F2 active site.

### Metabolic diversification of N-acyl amino acids downstream of CYP4F2

CYP4F2-catalyzed substrate hydroxylation can result in either metabolic diversification (e.g., arachidonic acid to 20-HETE) or oxidative degradation (e.g., vitamin E). To determine which of these metabolic outcomes was the relevant pathway for the N-acyl amino acids, we used untargeted lipidomics in live cells to map the fate of N-oleoyl-leucine and N-oleoyl-phenylalanine downstream of CYP4F2. For these studies, we initially focused on N-oleoyl-leucine. First, CYP4F2 or mock transfected HEK293T cells were treated with N-oleoyl-leucine for 4 h. Total intracellular lipids were extracted by the Folch method and differential peaks were identified using XCMS^22^ (**Fig. 5A**). Two statistically significant peaks of high fold change (P<0.05, >20-fold) were enriched in cells overexpressing CYP4F2. As expected, the top scoring peak of m/z = 410.3271 matched to ω-hydroxy-oleoyl-leucine (expected m/z = 410.3276) (**Fig. 5B**). The second peak of mass m/z = 674.5718 was also highly enriched in *CYP4F2*-transfected cells, but its chemical structure remained initially unknown. The mass difference observed between the two peaks (264.2442) as well as the later retention time of the second peak was consistent with an additional lipidation by oleate (+C_18_H_32_O, expected +264.2453). We therefore hypothesized that this second peak represented an unusual and previously unknown very long chain fatty acid, oleic acid-hydroxy-oleoyl-leucine (OAHOL). We used chemical synthesis to generate authentic standards for both hydroxy-oleoyl-leucine and OAHOL (see **Methods**). As shown in **Fig. 5C-D**, fragmentation of both synthetic hydroxy-oleoyl-leucine and the endogenous m/z = 410.33 peak gave an identical daughter ion at m/z = 130.087, which matched the mass of the leucine anion. In addition to demonstrating the existence of these metabolites in cell lysates we confirmed the presence of endogenous hydroxy-oleoyl-leucine in human plasma as well (**Fig. 5C, lower panel**). Similarly, fragmentation of both synthetic OAHOL and the endogenous m/z = 674.57 mass gave daughter ions at m/z = 130.087 (leucine anion) and m/z = 281.249 (oleate). We therefore conclude that m/z = 410.33 and m/z = 674.57 correspond to hydroxy-oleoyl-leucine and OAHOL, respectively.

**Fig. 5.**
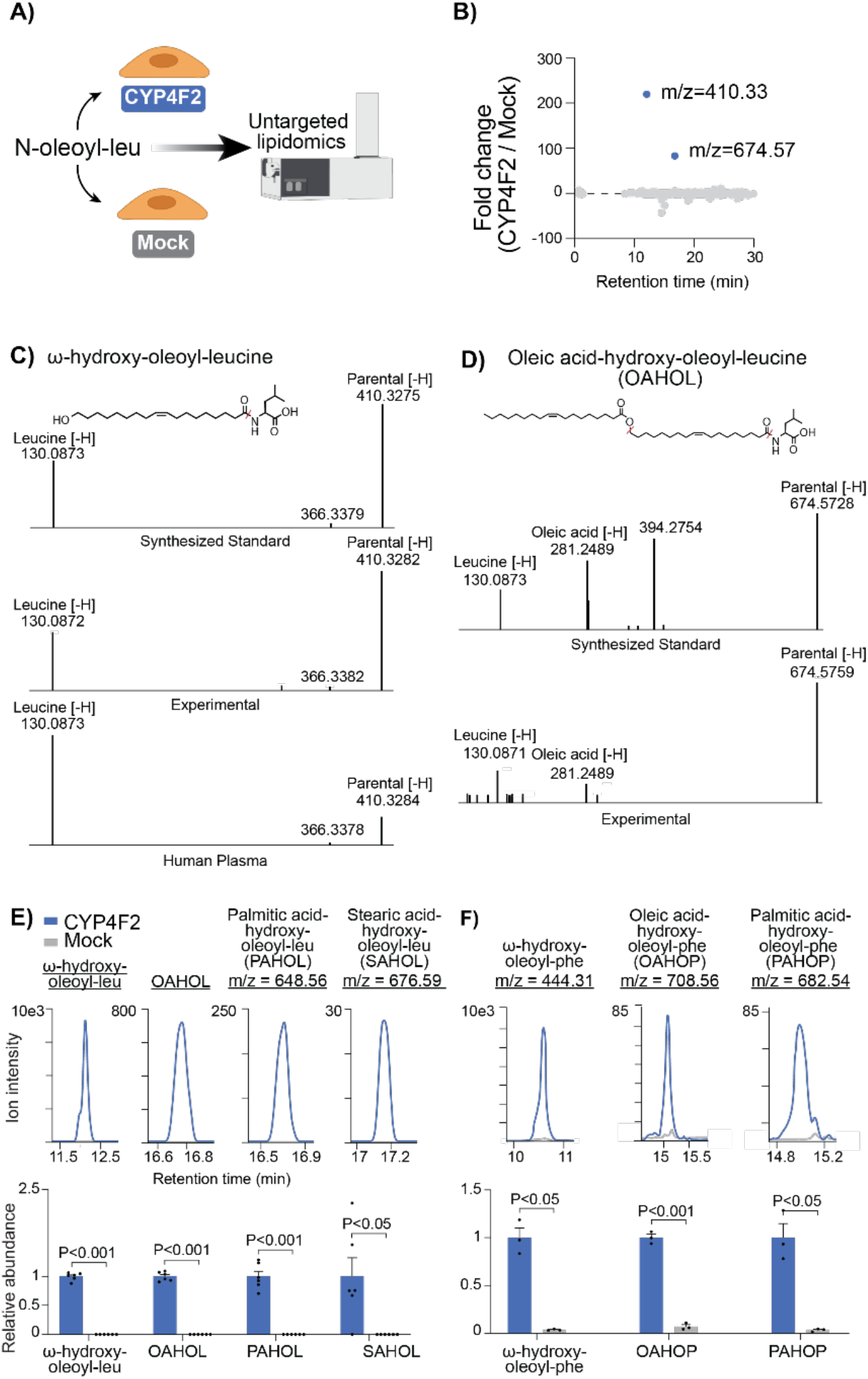
Untargeted lipidomics reveals very long chain N-oleoyl-leucine/phenylalanine derivatives downstream of CYP4F2. (A) Schematic of N-oleoyl-leucine live cell tracing experiment performed in CYP4F2 and mock transfected cells. (B) Plot of all significantly changed lipids (p < .05) detected by untargeted lipidomics in CYP4F2 versus mock transfected HEK293Ts treated with N-oleoyl-leucine (N=6). (C) Structural assignment and tandem MS fragmentation of an ω-hydroxy-oleoyl-leucine standard (top), experimental *m/z*=410.3282 mass (middle), and endogenous in human plasma *m/z*=410.3284 mass (bottom). (D) Structural assignment and tandem MS fragmentation of an OAHOL standard (top) and experimental *m/z*=674.5759 mass (bottom). The m/z=394.2754 mass is likely a background MS2 ion contaminant. As shown in Supplemental Fig. 2, the extracted ion chromatogram for 674.5728>394.2754 does not co-elute with the parent 674.5728, and also does not co-elute with bona fide transitions (e.g., the leucine transition 674.5728>130.0873). All other unannotated peaks are low abundance background. (E) Representative extracted ion chromatograms and abundance of ω-hydroxy-oleoyl-leucine, OAHOL, PAHOL, and SAHOL detected in CYP4F2 versus mock transfected cells (N=6). (F) Representative extracted ion chromatograms and abundance of ω-hydroxy-oleoyl-phenylalanine, OAHOP, and PAHOP detected in CYP4F2 versus mock transfected cells (N=3). Data are shown as mean ± SEM.

OAHOL is a novel lipid metabolite that is not found in any public databases^23, 24^. This lipid metabolite is presumably formed by enzymatic acylation of hydroxy-oleoyl-leucine with oleoyl-CoAs. Based on this metabolic pathway hypothesis, we manually examined our dataset for additional fatty acyl-derivatives of ω-hydroxy-oleoyl-leucine. Beyond OAHOL, we also identified PAHOL (palmitic acid-hydroxy-oleoyl-leucine) and SAHOL (stearic acid-hydroxy-oleoyl-leucine) dramatically elevated in CYP4F2-transfected, but not mock-transfected cells (**Fig. 5E** **and Supplemental Fig. 3A-B**). To determine if these very long chain lipid derivatives could also be formed downstream of CYP4F2-mediated hydroxylation of N-oleoyl-phenylalanine, we performed similar untargeted experiments in live CYP4F2-transfected cells with N-oleoyl-phenylalanine as a substrate. As shown in **Fig. 5F**, addition of N-oleoyl-phenylalanine also resulted in the downstream production of ω-hydroxy-oleoyl-phenylalanine and oleoyl- and palmitoyl-conjugates of hydroxy-oleoyl-phenylalanine (oleic acid-hydroxy-oleoyl-phenylalanine and palmitic acid-hydroxy-oleoyl-phenylalanine, respectively). For N-oleoyl-phenylalanine, the stearoyl acylation product was not detected. Taken together, we conclude that CYP4F2-catalyzed hydroxylation of N-acyl amino acids leads to the metabolic diversification of a wide array of previously unknown lipids, including several very long chain lipids that structurally resemble previously reported anti-diabetic fatty acid hydroxy fatty acids (FAHFAs^14^, **Supplemental Fig. 3C**).

## Discussion

By integrating genetic and clinical data with circulating levels of plasma N-acyl amino acids from a large human cohort, our study provides several insights of potential importance into the regulation of N-acyl amino acids in humans. First, individual plasma N-acyl amino acid levels are associated with cardiometabolic disease endpoints independent of free amino acid plasma levels and in patterns according to the amino acid head group. Second, the underlying genetic architecture and biological pathways that may regulate plasma levels of N-acyl amino acids also differ according to the amino acid head group. Finally, CYP4F2 functions as a human-specific enzyme node that catalyzes the metabolic diversification of N-acyl amino acids into a much larger family of lipids with varying characteristics of the fatty acid tail group, including several that structurally resemble FAHFAs.

Our large-scale human study revealed novel associations between the N-acyl amino acids and specific cardiometabolic diseases in participants of the JHS. Notably, we identified that the associations between fasting baseline levels of N-oleoyl-serine and future coronary heart disease, as well as N-oleoyl-leucine and future heart failure, were independent of the plasma levels of free serine and leucine, respectively. This suggests that these associations are driven by the intact N-acyl oleic acid conjugate of each amino acid, rather than levels of the free amino acid themselves. This may suggest that N-acyl amino acids affect cardiometabolic outcomes through distinct biological mechanisms from those mediated by circulating levels of the corresponding free amino acid.

Interestingly, when analyzing associations of N-acyl-amino acids with prevalent cardiometabolic disease at baseline, we detected a bifurcation in associations according to amino acid head group, with N-oleoyl-glycine and serine associated with cardiometabolically “advantageous” traits (such as prevalent diabetes), and N-oleoyl-leucine and phenylalanine associated with “disadvantageous” cardiometabolic traits (such as obesity). This division mirrors positive associations between branched chain (e.g. leucine) and aromatic (e.g. phenylalanine) free amino acids with obesity and insulin resistance, and inverse associations between glycine and serine with impaired glucose tolerance and risk of diabetes^25–33^. The mechanisms linking these free amino acid species to cardiometabolic traits and risk of disease remain to be fully elucidated, but may involve various opposing effects on regulatory pathways upstream of pancreatic islet β-cell insulin secretion^25, 34–38^. Whether the N-oleoyl species of these amino acids – as well as the CYP4F2-mediated derivatives of these metabolites – act through the same or different pathways adds a potential additional layer of complexity to this regulatory balance.

Our studies of human N-acyl amino acid metabolism highlight the utility of integrating metabolomic profiling data with genetic data for pathway discovery. In mice, FAAH and PM20D1 are two major enzymes that catalyze the bidirectional N-acyl amino acid synthesis and hydrolysis from free amino acids and free fatty acids. Human-based studies have established FAAH as the primary degradative enzyme of the structurally-related N-acyl ethanolamines^39^. Therefore, the association between the *FAAH* locus and plasma levels of N-oleoyl-glycine and N-oleoyl-serine in participants of the JHS was expected. However, the absence of an association of N-acyl amino acids to the human *PM20D1* locus, as well as the identification of the *CYP4F2* gene as a novel and human-specific association, were both unexpected. We validated the novel genetic association between N-oleoyl-leucine and N-oleoyl-phenylalanine with the *CYP4F2* locus by demonstrating that these two N-acyl amino acids are both bona fide substrates and inhibitors of the human CYP4F2 enzyme in vitro and in cell-based models. The top association for both N-oleoyl-leucine and N-oleoyl-phenylalanine was with a missense variant (chr19:15879621C>T; Val433Met) that has previously been shown to result in reduced CYP4F2 protein levels and enzymatic activity in cell-based model systems, but that has not previously been tied to N-acyl amino acid metabolism^20, 40, 41^. This variant is common with a global MAF of 0.29 and a subgroup MAF of 0.09 in African Americans, as reported in the NCBI Allele Frequency Aggregator.

While it is possible that our study did not detect an association between the *PM20D1* locus and N-acyl amino acids due to cohort-specific characteristics or sample size, we note that none of the measured N-acyl amino acids demonstrated even a modest association (nominal p-value ≤ 0.001) with a genetic variant between the *PM20D1* transcriptional start and end sites. This finding will require validation in additional cohorts, however may suggest that human PM20D1 controls local, and possibly tissue-specific extracellular paracrine pools of N-acyl amino acids that do not interact with the levels found in the circulation.

It is notable that we detected very strong associations between the *CYP4F2* locus on chromosome 19 and N-oleoyl-leucine and N-oleoyl-phenylalanine with no appreciable association signals between this locus and either N-oleoyl-glycine or N-oleoyl-serine (p > 0.001 for all variants between the *CYP4F2* transcriptional start and end sites). Importantly, the biochemical basis for this genetic specificity remains the subject of future study. We have previously found that subcellular location, and even organ-specific substrate accessibility can contribute to the regulation of specific N-acyl amino acids by FAAH and PM20D1^4^. Future detailed enzymology studies of CYP4F2 may uncover additional fundamental biochemical characteristics of this enzyme that underlie these specific genetic associations.

Our data suggest that CYP4F2 appears to be a key enzymatic node at the center of a complex lipid network that contains classical bioactive lipids, such as arachidonate and 20-HETE, with more novel lipid species, including N-acyl amino acids, hydroxylated N-acyl amino acids, and very long chain fatty acyl derivatives of N-acyl amino acids that structurally resemble FAHFAs. Notably, we detect that examples of these, including hydroxylated oleoyl-leucine, are present in human plasma. Currently, the precise biochemical and functional relationship between all of these lipid species, especially in a complex tissue environment such as a human liver, remains unknown. For instance, it may be possible that hydroxy N-acyl amino acids exhibit similar vasoactive effects as 20-HETE, and fatty acylated hydroxy-N-acyl amino acid derivatives might also function in a similar manner to that of the anti-diabetic FAHFAs. Alternatively, N-acyl amino acids may indirectly modulate the levels of other CYP4F2-regulated lipids via substrate competition. An important area of future work will be to understand the relative fluxes of each lipid class through CYP4F2 as well as the potential functional roles of these new N-acyl amino acid derivatives.

The integration of human functional genomics with untargeted lipidomics studies of cultured cells further allowed for the discovery of a novel class of very long chain lipids. These compounds structurally resemble FAHFAs, although instead of containing a characteristic branched hydroxyl linkage, contain a terminal ester linkage between a fatty acid and a hydroxy N-acyl amino acid. A similar terminal linkage has been described in O-acyl-ω-hydroxy fatty acids (ωOAHFAs), however these unusual species have only recently been detected in human skin^42^ and meibum^43^, and not to our knowledge in human plasma. Further, we are not aware of a previous report of fatty acyl-derivatives of hydroxy N-acyl-amino acids. FAHFAs are a diverse family of bioactive lipids that have recently been shown to signal through specific G protein-coupled receptors and other pathways to improve glucose-insulin homeostasis and block inflammatory cytokine production^14^. Our data provide new insights to this emerging class of molecules and raise the intriguing possibility that ω-hydroxy-oleoyl amino acid species may also serve as signaling effectors in human metabolism.

Finally, we note that the *CYP4F2* locus has been associated with several cardiometabolic phenotypes in large-scale, consortium-based GWAS meta-analyses, including blood total cholesterol levels (z-stat 5.24, p-value=8.1 x 10^-8^), LDL cholesterol levels (z-stat 4.77, p-value=9.2 x 10^-7^), diastolic blood pressure levels (z-stat 3.75, p-value=8.8 x 10^-5^), and coronary artery disease in subjects without diabetes (z-stat 3.28, p-value=5.3 x 10^-4^)^44^. Notably, the rs2108622 variant, which results in the CYP4F2 V433M substitution that abrogates CYP4F2 hydroxylation activity (**Fig. 3D-E**), is associated with several cardiometabolic phenotypes, including diastolic blood pressure (beta 0.008, p-value=1.13×10-7), total cholesterol (beta 0.006, p-value=9.54×10-7), Non-HDL cholesterol (beta 0.008, p-value=4.71×10-6), and coronary artery disease (OR 1.02, p=1.92×10-5)^45^. How this locus contributes to these human phenotypes remains the subject of future investigations, and the link with N-acyl amino acid biology may provide new insight to these studies.

## Methods

### Human Cohort

JHS is a prospective population based observational study designed to investigate risk factors for cardiovascular disease (CVD) in Black individuals, as previously described^46^. In 2000-2004, 5306 Black individuals from the Jackson, Mississippi tri-county area (Hinds, Rankin and Madison counties) were recruited for a baseline examination. Of the original cohort, 2351 individuals had metabolomic profiling of N-acyl amino acids performed from baseline samples and were included in the analyses. Details in regard to the collection and calculation of clinical data included in Supplemental Table 1 and Figure 1 have been previously described^13^. All clinical data were collected during the baseline exam (Exam 1), except visceral adipose tissue (Exam 2), subcutaneous adipose tissue (Exam 2), coronary artery calcium score (Exam 2), and abdominal aorto-iliac calcium score (Exam 2).

### Study Approval

The Institutional Review Boards of Beth Israel Deaconess Medical Center and the University of Mississippi Medical Center approved the human study protocols, and all participants provided written informed consent.

### N-Acyl amino acid metabolite profiling in human plasma

Methods for metabolomics profiling in human plasma have been described^12^. In brief, to measure N-oleoyl-leucine/phenylalanine/glycine/and serine, chromatography was performed using an Agilent 1290 infinity LC system equipped with a Waters XBridge Amide column, coupled to an Agilent 6490 triple quadrupole mass spectrometer. To measure endogenous hydroxy-oleoyl-leucine, chromatography was performed using a Waters UPLC BEH Amide (1.7um, 1.0×150mm) column. Metabolite transitions were assayed using a dynamic multiple reaction monitoring system. LC-MS data were analyzed with Agilent Masshunter QQQ Quantitative analysis software. Isotope labeled internal standards were monitored in each sample to ensure proper MS sensitivity for quality control. Pooled plasma samples were interspersed at intervals of ten participant samples to enable correction of drift in instrument sensitivity over time and to scale data between batches. A linear scaling approach was used to the nearest pooled plasma sample in the queue.

### Genotyping

Whole genome sequencing (WGS) in JHS has been described^47^. Briefly, participant samples underwent >30× WGS through the Trans-Omics for Precision Medicine project at the Northwest Genome Center at University of Washington and joint genotype calling with participants in Freeze 6; genotype calling was performed by the Informatics Resource Center at the University of Michigan.

### Whole Genome Association Studies

Summary statistics from whole genome association studies of metabolite levels in plasma samples of participants of the JHS are available on the NHGRI-EBI GWAS Catalogue (Accession GCP000239)^15^ and GWAS methods have been previously described^16^. Briefly, metabolite LC-MS peak areas were log-transformed and scaled to a mean of zero and standard deviation of 1 and subsequently residualized on age, sex, batch, and principal components (PCs) of ancestry 1-10 as determined by the GENetic EStimation and Inference in Structured samples (GENESIS)^48^, and subsequently inverse normalized. The association between these values and genetic variants was tested using linear mixed effects models adjusted for age, sex, the genetic relationship matrix, and PCs 1-10 using the fastGWA model implemented in the GCTA software package^49^. Variants with a minor allele count less than 5 in JHS were excluded from analysis.

### Cell line cultures

All cell lines were grown at 37°C with 5% CO2. HEK293T cells were obtained from ATCC and grown in Dulbecco’s modified Eagle’s medium (DMEM) with 10% fetal bovine serum (FBS) and 1% penicillin/streptomycin (pen/strep).

### Production of recombinant enzymes

Human CYP4F2-myc-DDK (OriGene RC216427), human

POR (OriGene SC100401), and human CYB5R1-myc-DDK (OriGene RC205833L3) plasmids were transiently co-transfected into HEK293T cells using PolyFect (Qiagen 301105) according to the manufacturer’s instructions. The medium was changed one day post-transfection. After an additional 24 hrs, the cells were harvested by scraping.

### Generation of CYP4F2 V433M mutant

Site-directed mutagenesis was performed using the Q5

Site-Directed Mutagenesis Kit (NEB E0554). Following the manufacturer’s instructions, PCR was performed using the human CYP4F2-myc-DDK (OriGene RC216427) plasmid (Forward primer, 5’-CCCTGAGGTCTACGACCC-3’; reverse primer, 5’-TCCGGCCACATAGCTGGGTTG-3’).

### Quantitative PCR analysis

Quantitative PCR (qPCR) was performed using SYBR Green qPCR

Master Mix (Bimake B21202) according to the manufacturer’s instructions. The following primers, written 5’ to 3’, were used for measuring the indicated genes: *TBP*, ACCCTTCACCAATGACTCCTATG and TGACTGCAGCAAATCGCTTGG; *CYP4F2*, GAGGGTAGTGCCTGTTTGGAT and CAGGAGGATCTCATGGTGTCTT.

Untargeted measurements of metabolites by LC-MS. Untargeted metabolomics measurements were performed on an Agilent 6520 Quadrupole Time-of-Flight (Q-TOF) LC/MS. Mass spectrometry analysis was performed using electrospray ionization (ESI) in negative mode. The dual ESI source parameters were set as follows, the gas temperature was set at 250 °C with a drying gas flow of 12 l/min and the nebulizer pressure at 20 psi. The capillary voltage was set to 3500 V and the fragmentor voltage set to 100 V. Separation of metabolites was conducted on a Luna 5 µm C5 100 Å, LC Column 100 x 4.6 mm (Phenomenex 00D-4043-E0) with normal phase chromatography. Mobile phases were as follows: Buffer A, 95:5 water/methanol; Buffer B, 60:35:5 isopropanol/methanol/water with 0.1% ammonium hydroxide in both Buffer A and B for negative ionization mode. For 10 minute runs, the LC gradient started at 95% A with a flow rate of 0.6 ml/min from 0-1 min. The gradient was then increased linearly to 95% B at a flow rate of 0.6 ml/min from 1-8 minutes. From 8-10 minutes the gradient was maintained at 95% A at a flow rate of 0.6 ml/min. For 30 minute runs, the LC gradient started at 95% A with a flow rate of 0.6 ml/min from 0-3 min. The gradient was then increased linearly to 95% B at a flow rate of 0.6 ml/min from 3-25 minutes. From 25-30 minutes the gradient was maintained at 95% A at a flow rate of 0.6 ml/min.

Targeted metabolomics in cell culture samples. Targeted measurements were performed on an Agilent 6470 Triple Quadrupole (QQQ) LC/MS. Mass spectrometry analysis was performed using electrospray ionization (ESI) in negative mode. The AJS ESI source parameters were set as follows, the gas temperature was set at 250 °C with a gas flow of 12 l/min and the nebulizer pressure at 25 psi. The sheath gas temperature was set to 300 °C with the sheath gas flow set at 12 l/min. The capillary voltage was set to 3500 V. Separation of metabolites was performed as described above in the untargeted metabolomics section. Multiple reaction monitoring (MRM) was performed for the indicated metabolites with the listed dwell times, fragmentor voltage, collision energies, cell accelerator voltages, and polarities.

| Compound | Precursor Ion | MS1 Res | Product Ion | MS2 Res | Dwell | Fragmentor | Collision Energy | Cell Accelerator Voltage | Polarity |
| --- | --- | --- | --- | --- | --- | --- | --- | --- | --- |
| N-oleoyl-leucine | 394.4 | Unit | 130.1 | Unit | 50 | 135 | 30 | 5 | Negative |
| ω-hydroxy-oleoyl-leucine | 410.3 | Unit | 130.1 | Unit | 50 | 135 | 30 | 5 | Negative |
| Arachidonic acid | 303.2 | Unit | 303.2 | Unit | 50 | 135 | 0 | 5 | Negative |
| 20-HETE | 319.2 | Unit | 319.2 | Unit | 50 | 135 | 0 | 5 | Negative |
| N-oleoyl-phenylalanine | 428.3 | Unit | 164.1 | Unit | 50 | 135 | 30 | 5 | Negative |
| ω-hydroxy-oleoyl-leucine | 444.3 | Unit | 164.1 | Unit | 50 | 135 | 30 | 5 | Negative |
| OAHOP | 708.6 | Unit | 164.1 | Unit | 50 | 135 | 30 | 5 | Negative |
| PAHOP | 682.5 | Unit | 164.1 | Unit | 50 | 135 | 30 | 5 | Negative |

### Synthesis of ω-hydroxy-oleoyl-leucine

1-Ethyl-3-(3-dimethylaminopropyl)carbodiimide hydrochloride (EDC-HCl, 1.05 eq) was added to a cold mixture (ice bath) of ω-hydroxy-oleic acid (A2B Chem BG26794, 1 eq) and 1-hydroxybenzo-triazole monohydrate (HOBt H2O, 2 eq) in N,N-dimethyformamide (DMF). Stirring was continued for 10 min before a mixture of L-leucine ethyl ester hydrochloride (5 eq) and N,N-diiso-propylethylamine (DIPEA, 3 eq) in DMF was added. The ice bath was removed, and the reaction mixture was stirred for 16 h at room temperature. After removal of the solvent, the residue was taken up in ethyl acetate (EtOAc); washed with KHSO4, brine, NaHCO3, and brine; and then concentrated under reduced pressure. The crude ester was dissolved in tetrahydrofuran, and 2 N of LiOH was added. The resulting mixture was stirred at room temperature for 4 h. The reaction was acidified to pH 2-3 by the addition of 1 M of HCl and then extracted with DCM and concentrated. The resulting crude product was used directly for LC-MS/MS analysis.

### Synthesis of oleic acid-hydroxy-oleoyl-leucine

To a solution of oleic acid in dichloromethane (DCM) was added oxalyl chloride (one drop) and one drop of DMF at 0 C. The mixture was stirred at room temperature for 2 hours. The mixture was concentrated and dissolved in DCM then added to a suspension of ω-hydroxy-oleoyl-leucine and DIPEA (1 drop) in DCM. The reaction mixture was stirred at room temperature for 16 hours then acidified to pH 4 with 1 M HCl. The resulting mixture was extracted with DCM and washed with brine then the solvent was removed under reduced pressure. The crude product was analyzed directly by LC-MS/MS.

### Lipid analysis by LC-MS/MS in cell culture samples

To acquire tandem mass (MS2) spectra for N-oleoyl-leucine and its derivatives, lipid samples were first dried by a SpeedVac and then reconstituted in acetonitrile:isopropanol:water mixture at a ratio of 13:6:1 (v/v/v). Samples were injected at 4µL into an Acsentis Express C18 2.7 µm HPLC column (L × I.D. 15 cm × 2.1 mm, Millipore Sigma 53825-U) preceded by an Acsentis Express C18 2.7 µm guard column (5 x 2.1 mm guard (L × I.D. 5 mm × 2.1 mm, Millipore Sigma 53501-U). Flow rate was set to a constant rate 0.26 mL/min. Additional chromatographic details are as follow:

Mobile phases: A, 10 mM ammonium formate and 0.1% formic acid in 60% and 40 water and acetonitrile, respectively; B, 10 mM ammonium formate and 0.1% formic acid in 90% and 10% 2-propanol and acetonitrile, respectively. Separation gradient: isocratic elution at 32% B from 0−1.5 min; linear increase from 32-45% B from 1.5-4 min; linear increase from 45-52% B from 4-5 min; linear increase from 52-58% B from 5-8 min; linear increase from 58-66% B from 8-11 min; linear increase from 66-70% B from 11-14 min; linear increase from 70-75% B from 14-18 min; linear increase from 75-97% B from 18-21 min; hold at 97% B from 21-35 min; linear decrease from 97-32% B from 35−35.1 min; hold at 32% B from 35.1-40 min.

An ID-X tribrid mass spectrometer (Thermo Fisher Scientific) with a heated electrospray ionization (HESI) probe was used for both full scan mass (MS1) and MS2 spectra acquisition. sheath gas, 40 units; aux gas, 10 units; sweep gas, 1 unit; ion transfer tube temperature, 300 ◦C; vaporizer temperature, 375 ◦C; negative ion voltage, 3000 V; orbitrap resolution MS1, 120,000, MS2, 30,000; scan range (m/z), 220-1500; RF lens, 40%; AGC target MS1, 4×10^5^, MS2, 5×10^4^; maximum injection time MS1, 50 ms, MS2, 54 ms; data-dependent tandem mass spectrometry (ddMS2) isolation window (m/z), 1; activation type, higher energy collision dissociation (HCD); HCD fragmentation, stepped 15%, 25%, 35%; cycle time, 1.5s; microscans, 1 unit; intensity threshold, 1.0e4; dynamic exclusion time, 2.5 s. A mass list corresponding to the [M-H]^-^ adducts of the parental lipids was used for targeted MS2 fragmentation. EASYICTM was enabled for internal calibration.

### *In vitro* CYP42 assays

HEK293T cells were co-transfected with CYP4F2 WT/V433M, POR, and CYB5R1. CYP4F2 and mock transfected cells were harvested in PBS, lysed by sonication, and centrifuged (10 min at 15,000 rpm) to remove debris. In vitro enzymatic reactions were conducted in 96-well plates and initiated with 1 mM NADPH. Final reaction conditions were 100 µM substrate (N-oleoyl-leucine, Cayman Chemical 20064, N-oleoyl-phenylalanine, Cayman Chemical 28921, or arachidonic acid, Cayman Chemical 90010) and 50 µg protein in 50 µl PBS. After 1 hr. at 37°C, reactions were transferred to glass vials, quenched with 150 µl 2:1 v/v chloroform:methanol, and vortexed. Reaction vials were centrifuged (10 min at 1000 rpm) and the organic layer was transferred to a mass spec vial and analyzed by LC-MS as described above. For competition assays, cell lysates were incubated for 5 min with competitor (Arachidonic acid, docosanoic acid, Cayman Chemical 9000338, or 8-HETE, Cayman Chemical 34340) at 37°C before addition of other substrates and initiation with NADPH.

### Kinetic enzymatic assays

CYP4F2 lysates were harvested from transiently transfected HEK293T cells as described above. In vitro enzymatic reactions were conducted in 96-well plates and initiated with 1 mM NADPH. Final reaction conditions were: 1 µM, 4 µM, 20 µM, and 100 µM of substrate and 0.46 µg CYP4F2 enzyme in 50 µl PBS. After 10 min at 37°C, reactions were transferred to glass vials quenched with 150 µl 2:1 v/v chloroform:methanol, and vortexed. Reaction vials were centrifuged (10 min at 1000 rpm) and the organic layer was transferred to a mass spec vial and analyzed by LC-MS as described above.

### Live cell tracing experiments

Transiently transfected (CYP4F2 or mock) HEK293Ts were washed twice with PBS and then harvested by scraping. Cells were spun down (5 min at 1000 rpm) and resuspended in serum free media then aliquoted into a 96-well plate (1.2 million cells per well). Final reaction conditions were 10 µM N-oleoyl-leucine or N-oleoyl-phenylalanine.

Reactions were initiated with 1 mM NADPH. After 4 hrs at 37°C, reactions were transferred to glass vials, quenched with 2:1 v/v chloroform:methanol, and vortexed. After centrifuging vials (10 min at 1,000 rpm), the organic layers were analyzed by LC-MS or LC-MS/MS as described above. LC-MS data were uploaded to Scripps XCMS Online to identify significantly changed metabolites.

### Western blot analysis

Cells were collected and lysed by sonication in PBS. Cell lysates were centrifuged at 4 °C for 10 minutes at 15,000 rpm to remove residual cell debris. Protein concentrations of the supernatant were normalized using the Pierce BCA protein assay kit and combined with 4 x NuPAGE LDS Sample Buffer with 10 mM DTT. Samples were then boiled for 10 minutes at 95 °C. Prepared samples were run on a NuPAGE 4-12% Bis-Tris gel then transferred to nitrocellulose membranes. Blots were blocked for 30 minutes at room temperature in Odyssey blocking buffer. Primary antibodies (mouse anti-FLAG and rabbit anti-Beta-actin) were added to Odyssey blocking buffer at a ratio of 1:1000. Blots were incubated in the indicated primary antibodies overnight while shaking at 4 °C. The following day, blots were washed 3 times with PBS-T, 10 minutes each before staining with the secondary antibody for 1 hour at room temperature. The secondary antibodies used were goat anti-rabbit and goat anti-mouse antibodies diluted in blocking buffer to a ratio of 1:10,000. Following secondary antibody staining, the blot was washed 3 times with PBS-T before being imaged with the Odyssey CLx Imaging System.

### Statistics

All data were expressed as mean ± SEM unless otherwise specified. A student’s t-test was used for pairwise comparisons. Unless otherwise specified, statistical significance was set as P < 0.05.

## Acknowledgements

Acknowledgement of funding sources

The Jackson Heart Study (JHS) is supported and conducted in collaboration with Jackson State University (HHSN268201800013I), Tougaloo College (HHSN268201800014I), the Mississippi State Department of Health (HHSN268201800015I) and the University of Mississippi Medical Center (HHSN268201800010I, HHSN268201800011I and HHSN268201800012I) contracts from the National Heart, Lung, and Blood Institute (NHLBI) and the National Institute on Minority Health and Health Disparities (NIMHD). The authors also wish to thank the staffs and participants of the JHS. Additional funding was provided from NHLBI K08HL145095 (to MDB), DK124265 (to JZL), NIH GM113854 (to VLL), the National Science Foundation (ALW), the Fundacion Alfonso Martin Escudero (to MDMG), NIH DP2-CA271389 and the Stanford Alzheimer’s Disease Research Center (to MAR), The Wu Tsai Human Performance Alliance at Stanford University and the Joe and Clara Tsai Foundation (to XL), and HHSN268201600034I, R01 NR019628, R01 DK081572, and the LeDucq Foundation (to REG).

## Disclaimer

The views expressed in this manuscript are those of the authors and do not necessarily represent the views of the National Heart, Lung, and Blood Institute; the National Institutes of Health; or the U.S.Department of Health and Human Services.

