## Supplemental Figures and Tables for "CYP4F2 is a human-specific determinant of circulating N-acyl amino acid levels"

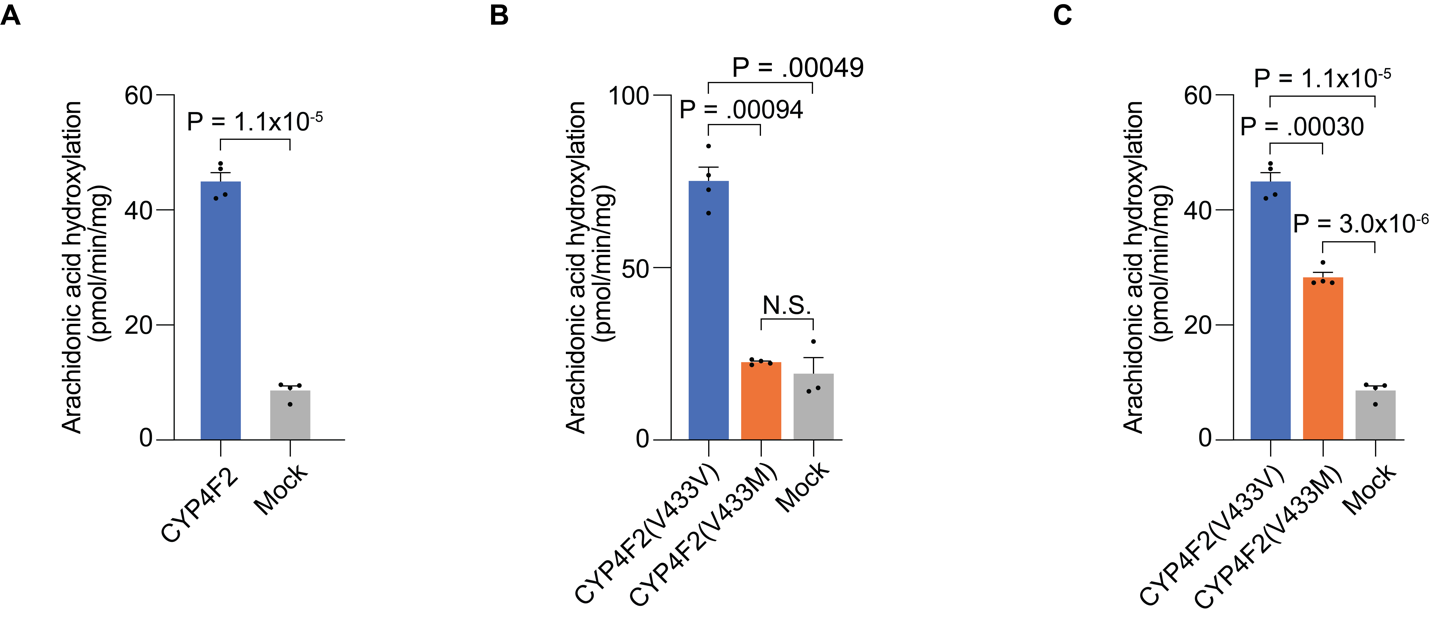

**Supp Fig. 1. Arachidonic acid is a canonical substrate of CYP4F2.**

(A) Arachidonic hydroxylation in cell lysates of CYP4F2-flag and mock transfected HEK293T cells (N=4). (B) Arachidonic acid hydroxylation in cell lysates of CYP4F2(V433V), CYP4F2(V433M), and mock transfected HEK293T cells (N=4). Cells were transfected with equal amounts of mRNA. (C) Arachidonic acid hydroxylation in cell lysates of CYP4F2(V433V), CYP4F2(V433M), and mock transfected HEK293T cells (N=4). Equal amounts of CYP4F2(V433V) and CYP4F2(V433M) enzyme were used.

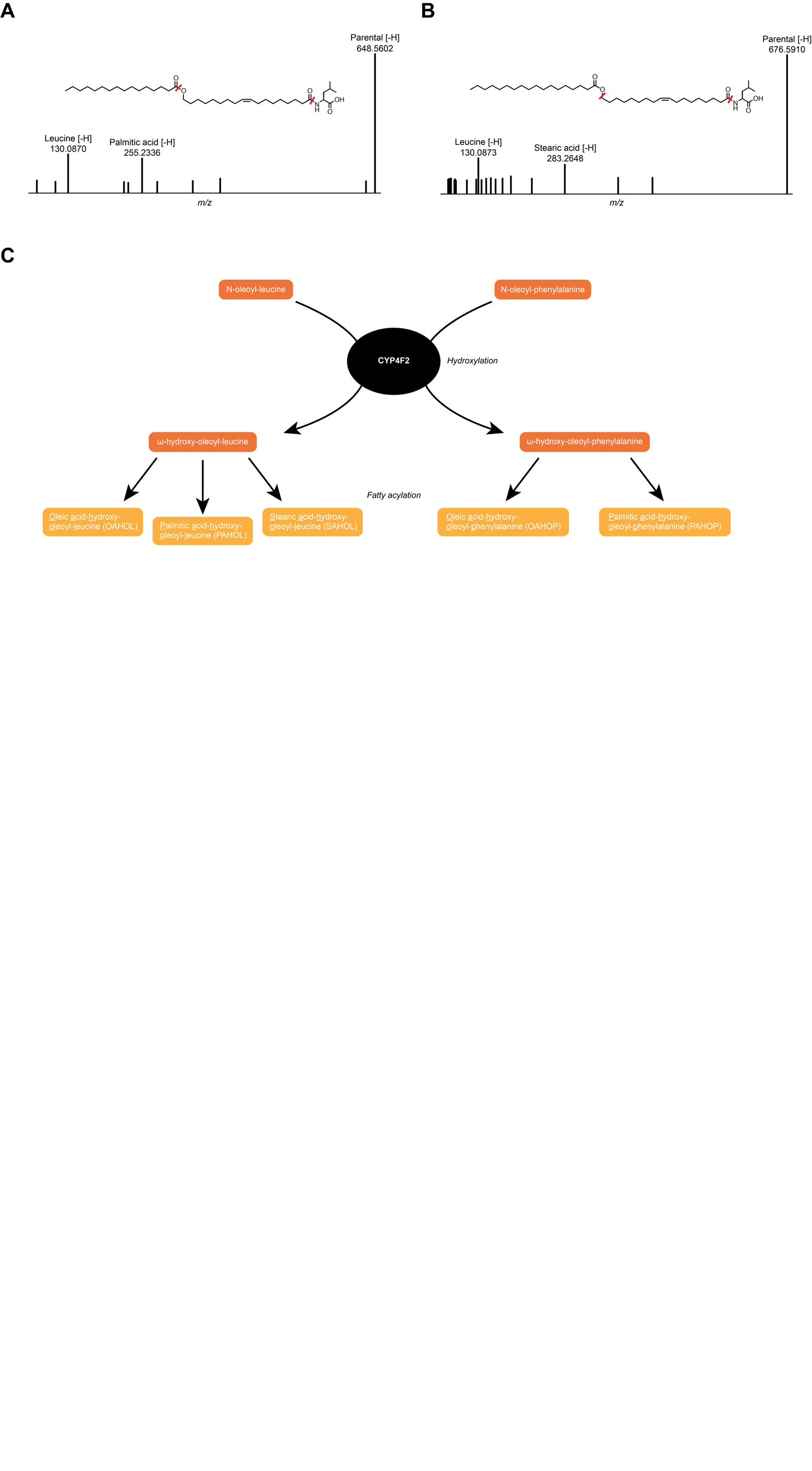

**Supp Fig. 2. Metabolic diversification of N-acyl amino acids downstream of CYP4F2**

(A) Tandem MS fragmentation of experimental *m/z*=648.5062 mass detected in CYP4F2 transfected cells treated with N-oleoyl-leucine. (B) Tandem MS fragmentation of experimental *m/z*=676.5910 mass detected in CYP4F2 transfected cells treated with N-oleoyl-leucine.

(C) Schematic demonstrating CYP4F2-mediated hydroxylation of N-oleoyl-leucine/phenylalanine and the subsequent formation of FAHOLs and FAHOPs.

**Supplementary Table 1.**

| **Clinical Characteristic** | **Full Cohort** | **N-Acyl-AAs Measured** |
| --- | --- | --- |
|  | **N = 5,306** | **N = 2,351** |
| Age, years | 55.4 (12.8) | 55.2 (12.6) |
| Male, n (%) | 1,934 (36%) | 899 (38%) |
| BMI, kg/m2 | 31.8 (7.2) | 31.8 (7.2) |
| eGFR, mL/min/1.73m2 | 85.7 (18.5) | 85.2 (18.9) |
| HbA1c, % | 6.0 (1.3) | 6.0 (1.4) |
| Fasting plasma glucose, mg/dL | 100.0 (33.4) | 101.3 (36.1) |
| Fasting plasma insulin, IU/mL | 18.2 (21.2) | 18.4 (25.1) |
| Systolic blood pressure, mmHg | 127.0 (18.4) | 127.0 (18.3) |
| Diastolic blood pressure, mmHg | 78.8 (10.6) | 79.1 (10.4) |
| Fasting total cholesterol, mg/dL | 199.3 (40.1) | 200.1 (40.4) |
| Fasting triglycerides, mg/dL | 106.4 (78.5) | 108.7 (73.5) |
| Fasting HDL cholesterol, mg/dL | 51.8 (14.6) | 51.6 (14.7) |
| Fasting LDL cholesterol, mg/dL | 126.6 (36.6) | 127.2 (37.3) |
| Prevalent diabetes mellitus, n (%) | 1,152 (22%) | 547 (23%) |
| Prevalent hypertension, n (%) | 3,252 (61%) | 1,452 (62%) |
| Prevalent coronary heart disease, n (%) | 400 (8%) | 145 (6%) |
| Statin medication use, n (%) | 602 (14%) | 261 (14%) |

Mean (SD); n (%)

Clinical characteristics were measured during visit 1 of the JHS

BMI: body mass index

eGFR: estimated glomerular filtration rate

HbA1c: Hemoglobin A1C

**Supplementary Table 2.**

| **Metabolite** | **Trait** | **Model 1 Estimate** | **Model 1 Std Error** | **Model 1 P-Value** | **Model 2 Estimate** | **Model 2 Std Error** | **Model 2 P-Value** |
| --- | --- | --- | --- | --- | --- | --- | --- |
| N-oleoyl-glycine | HbA1c | -0.17 | 0.02 | 7.72E-17 | -0.17 | 0.02 | 5.10E-16 |
| N-oleoyl-glycine | Fasting Insulin | -0.12 | 0.02 | 1.91E-07 | -0.12 | 0.02 | 3.55E-07 |
| N-oleoyl-glycine | HOMA-IR | -0.13 | 0.02 | 1.14E-08 | -0.13 | 0.02 | 2.43E-08 |
| N-oleoyl-glycine | Fasting Triglycerides | -0.14 | 0.02 | 1.56E-09 | -0.13 | 0.02 | 7.66E-09 |
| N-oleoyl-serine | HbA1c | -0.10 | 0.02 | 1.11E-06 | -0.11 | 0.02 | 3.68E-08 |
| N-oleoyl-serine | Fasting Insulin | -0.08 | 0.02 | 6.41E-04 | -0.09 | 0.02 | 4.82E-05 |
| N-oleoyl-serine | HOMA-IR | -0.08 | 0.02 | 1.42E-04 | -0.10 | 0.02 | 4.98E-06 |
| N-oleoyl-serine | Fasting Triglycerides | -0.15 | 0.02 | 3.37E-12 | -0.16 | 0.02 | 8.99E-14 |

Model 1: age, sex, metabolite batch-adjusted

Model 2: age, sex, metabolite batch, BMI-adjusted
